# Biohybrid toggle switches: leveraging bistability in muscle-actuated machines

**DOI:** 10.64898/2026.10.08.757177

**Authors:** Ronald H. Heisser, Ozgun K. Afsar, Pria Sawhney, Nicolas Castro, Inkyu Lee, Annika Marschner, Ritu Raman

## Abstract

Tissue-engineered skeletal muscle offers robotics unique advantages in energy density, adaptability, and self-repair, but current biohybrid machines couple muscle directly to artificial skeletons, limiting net force output and requiring continuous stimulation maintain mechanism positions. Here, we introduce a biohybrid bistable switching mechanism that converts antagonistic muscle contractions to snap-through actuation of an elastomeric four-bar flexure, maintaining stable positions without energy consumption. We decouple passive muscle pre-tensioning from flexure operation with thin-film elastomer “resistance bands,” enabling fine tuning of spring stiffness and force to match muscle mechanical output envelopes. Leveraging mesoscale additive manufacturing, spin coating, and replica molding techniques, components are integrated into a millimeter-scale linear stage to axially align muscles with artificial components. Our prototype achieves reversible toggling between bistable states with a 130 ms switching time, establishing skeletal muscle actuators as viable electromechanical switches, and broadening biohybrid robotics towards novel applications such as grippers, valves, and mechanical computers.

## Introduction

Tissue engineered skeletal muscle has potential to revolutionize robotics by merging artificial structures and mechanisms with the biological advantages of living tissue^1,2^. Muscle’s protein-based hierarchical architecture and volumetric chemical energy storage endow animals with high dexterity, strength, and energy density to navigate the outside world for durations far longer than that of motor-driven robots. Unlike synthetic actuators, muscle also grows, remodels, and repairs itself in response to injury, adapting to ever-changing environmental conditions. The microscale architecture of muscle sarcomeres sets specific tension (∼250 kPa in humans) and contractility (∼40% strain) that remain consistent across tissue length scales, resulting in the inspiring diversity of scale of motion seen across the animal kingdom^3^.

Recent research in micro-robotics has revealed a new domain of applications that favor multi-degree of freedom (DoF) miniature electromechanical systems to control miniature grippers^4^, morph surfaces^5^, optically steer lasers^6^, and power winged insect-scale robotic flight^7^. Investigations into millimeter-scale arrayed actuation systems have spawned new concepts for mechanical computation and fluidic logic that could be critical to advancing soft robotic dexterity and control^8^. The numerous actuation strategies employed to produce these demonstrations have power requirements and external equipment needs that limit device integration at the millimeter scale and the ability to achieve higher actuator packing densities^9,10^. Muscle naturally solves many of these challenges by storing and converting energy inside the actuator as ATP and glycogen, requiring low-power electrical signals to stimulate contractions that compactly transmit forces to external structures. Practically realizing muscle’s advantageous design with tissue engineered constructs requires developing compact, millimeter-scale biohybrid mechanisms that muscle can precisely actuate.

Current biohybrid robotics research focuses on integrating mm-scale engineered muscle into abiotic spring-like skeletons that enable rhythmic actuation and functional demonstrations of swimming^11^, crawling^12^, and pumping^13^ (Fig. 1A). In these designs, muscle cells anchored to a hydrogel extracellular matrix, represented by k_m_, need an artificial skeleton (k_s_) to keep them taut, as they would otherwise collapse their own shape and lose contractility^14^. Such designs come with two significant drawbacks. First, k_s_ has to be large enough to withstand the mN-scale passive tension forces generated by mm-scale muscles at rest, and thus limits active stroke in response to μN-scale forces generated during muscle contraction. Second, while spring-like skeletons are useful for rhythmic actuation, maintaining skeletons at a given position requires continuous muscle stimulation, leading to rapid muscle fatigue. These limitations point towards a critical need for biohybrid mechanisms that can decouple pre-tensioning mechanisms from actuation mechanisms, and can maintain actuation states without continuous power consumption. Accomplishing these goals would enable new applications for biohybrid robots that only require energy to switch between stable states such as valves, grippers, and mechanical computers (Fig. 1B).

**Figure 1:**
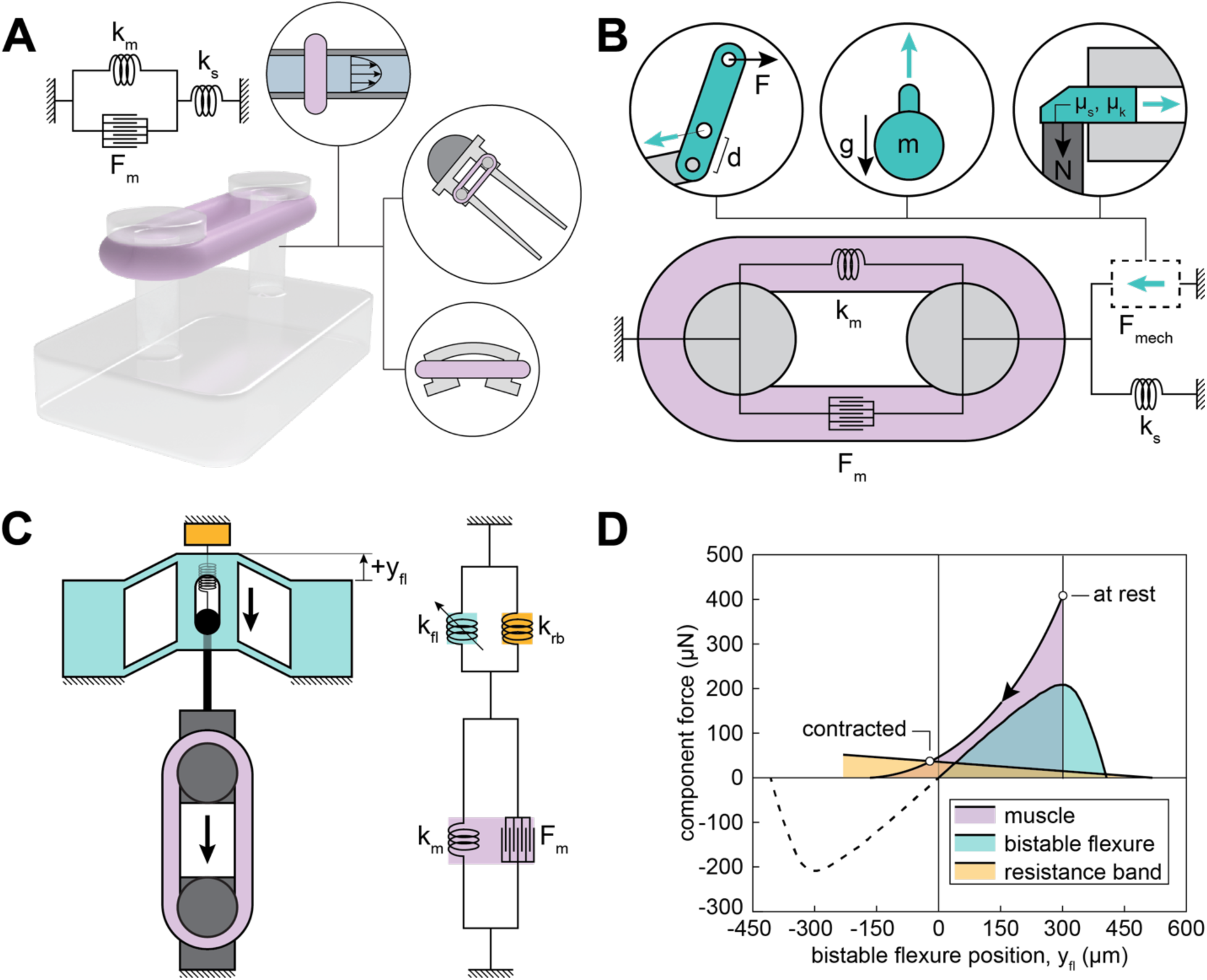
Biohybrid toggle system overview. A) The current biohybrid actuation paradigm couples passive muscle tensioning and functional mechanisms in a single abiotic spring-like structure, where muscle contraction deforms the structure to perform functions such as walking, swimming, or pumping. B) Designing within the mechanical output envelope of biohybrid actuators, additional mechanical components can be integrated with the muscles to yield more complex mechanisms that decouple passive tensioning and contraction. This design change enables pursuing designs beyond simple springs, such as joints, payloads, or latches^29^. The diagram in C) illustrates our proposed bistable actuation system, whereby a muscle is integrated with a soft pre-buckled flexure (k_fl_) and tensioning resistance band (k_rb_), flipping the flexure between two stable states when stimulated. The resistance band maintains homeostatic muscle tension even after the muscle loses engagement with the flexure. D) Flexure and tensioning spring stiffness and preloading can be tuned to fit under the muscle contractile force window, shown in pink.

To address the challenges limited with existing muscle-skeleton designs, we introduce the first example of a biohybrid bistable switching mechanism. This device uses antagonistic muscle contractions to drive snap-through of a buckled elastomeric four-bar flexure between two mechanically stable positions, consuming no energy while stationary (Fig. 1C). The simplest bistable mechanism would feature a muscle that is directly attached to a bistable flexure, where it is passively tensioned while the flexure remains upward. However, upon contraction and snap-through of the flexure, the muscle would then become slack and begin to collapse itself until the flexure reset upwards. Therefore, we decoupled passive muscle tensioning and mechanism function by including a “resistance band” elastomeric spring that restores the muscle to a stretched resting state, regardless of engagement with the flexure. Doing so allowed us to independently design component geometries and stiffnesses to match muscle actuator work output. We integrated our biohybrid machine components into a millimeter-scale linear stage and demonstrated reversible bistable actuation, establishing muscle’s function as an electromechanical switch.

## Results

### Biohybrid toggle switch design rationale

The force output of engineered skeletal muscles partly depends on the cell line. Actuators composed of primary and induced pluripotent stem cells (iPSCs) have demonstrated stress outputs of up to 50 kPa, yet each have their own limitations^3^. Primary cells require first extracting tissue from living mammals, iPSCs require sophisticated differentiation techniques that can take weeks. This article focuses on tissues built from immortalized C2C12 mouse myoblasts because of their potential to scale sustainably to large cell counts with minimal upstream biological processing. We engineered optogenetically modified C2C12 muscle actuators expressing a light-sensitive membrane calcium ion channels using previously established protocols to enable optical control of contractions, thus eliminating the need for integrated electrode design^15,16^. Currently, mm-scale C2C12 actuators generate stresses under 10 kPa, corresponding to forces up to ∼500 μN (blocked forces can be above 5 mN) and contractile strains under 10%^11,17,18^. To be conservative, we set mechanism design specifications to fit a 9 mm-long, 3 mm-wide actuator with a 5% strain (∼500 μm), 500 μN muscle contraction force envelope.

We previously coupled C2C12-derived muscles with flexures to measure contractile forces, finding that stiffnesses at or below 0.5 N/m maximized muscle stroke^14,19^. These flexures had planar dimensions on the order of 1000 mm^2^, two orders of magnitude larger than the muscle actuators themselves. Designing precise elastic components compatible with both muscle size and muscle force output requires designing with both soft materials and sub mm-features. We accomplished this by combining a series of mesoscale (∼10 μm to 1 mm) manufacturing processes to fabricate flexures that repeatably undergo snap-through displacements of under 500 μm and can be integrated into assemblies with millimeter-scale tensioning springs with under 0.5 N/m stiffnesses.

### Design and fabrication of resistance bands for muscle pre-tensioning

Muscles require passive tensioning elements that provide a static force to keep them taut. Following from Hooke’s law, *F* = *kx*, a particular spring force can be achieved by precisely picking spring stiffnesses and displacements. Within this design choice is an inherent tradeoff as, during contraction, the tensioning element stores elastic energy that muscle could apply to the desired actuation mechanism. Therefore, we designed towards minimizing stiffness, *k*, achieving requisite pre-tensioning forces with larger extensional displacements, *x*, that are difficult to implement with rigid materials without significantly increasing footprint. Inspired by sheet-style resistance bands used in physical therapy and exercise routines, we fabricated millimeter-scale resistance bands to efficiently provide muscle passive tension force.

Reaching a 0.5 N/m stiffness in a prismatic linear extension spring is not trivial, requiring at least one microscale cross-sectional dimension. The linear elastic axial stiffness of a rectangular beam is

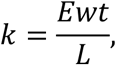

where *E* is the elastic modulus, *w* is the width, *t* is the thickness, and *L* is the length. Restricting spring resting length to 2.5 mm and width to 0.8 mm (matching the footprint of our muscle actuators), a millimeter-scale silicone (*E* = 100 kPa) resistance band would need a thickness of 15 μm to achieve a stiffness of 0.5 N/m. Therefore, we chose to spin coat silicone (Ecoflex 00-10, 100% modulus = 55 kPa) thin films in pursuit of sub-100 μm thicknesses (Fig. 2A)^20^. We characterized film thickness for a range of spin speeds and evaluated the impact of thinning additives. Adding thinner beyond 10% wt/wt caused films to lose their constitution and tear during substrate delamination. Spin coating at 3000 rpm or higher produced desired film thicknesses between 20 μm and 40 μm.

**Figure 2:**
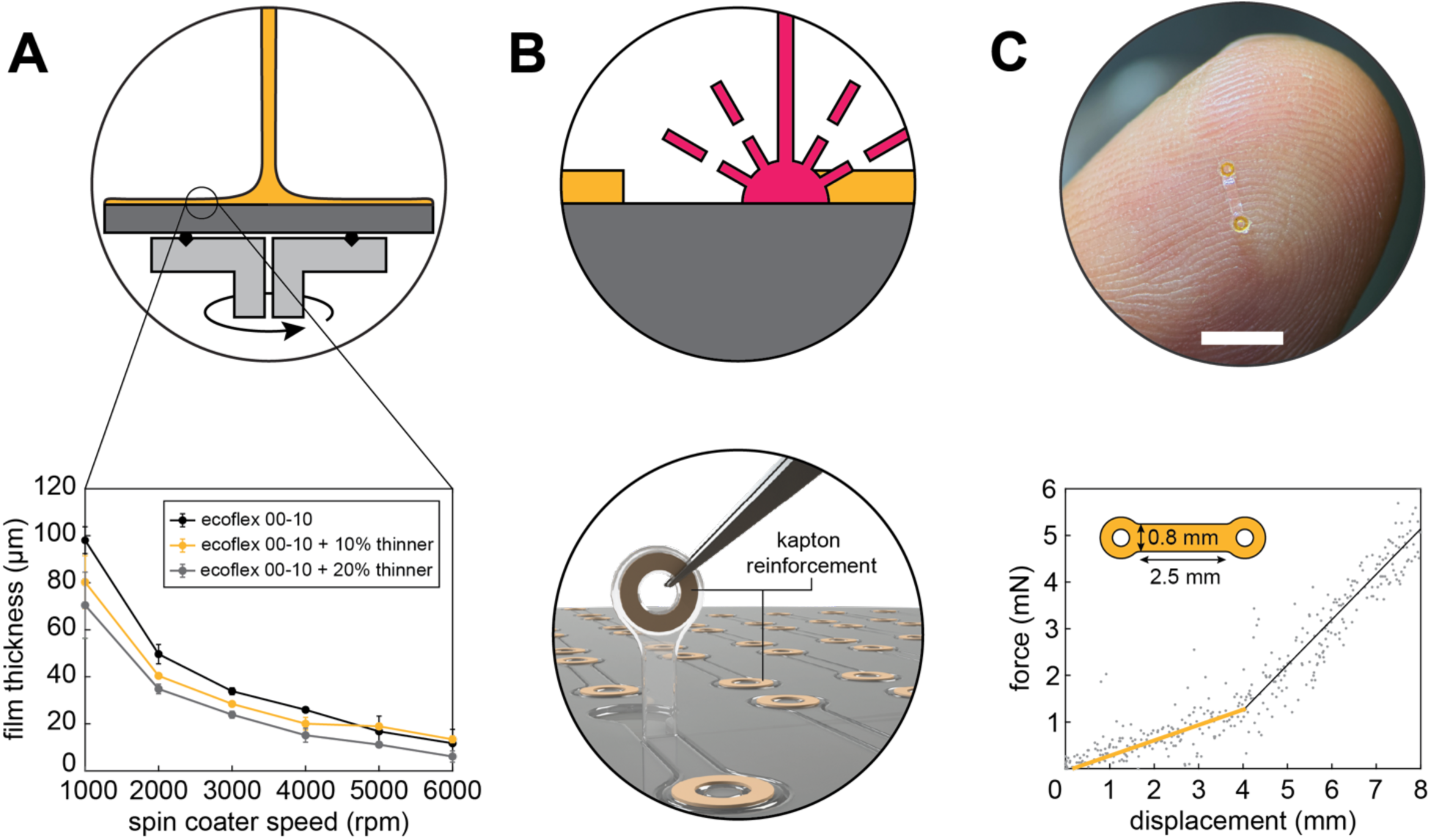
Resistance band manufacturing. We fabricated thin (<50 μm) silicone elastomer films that have stiffnesses under 0.5 N/m after being cut to the desired geometry. A) Film thickness was characterized for different formulations of Ecoflex 00-10 silicone thinner (Silicone Thinner, Smooth-On) mixtures, spin coating samples up to 6000 rpm. Film height was determined using a laser microscope. B) Resistance band patterns were created with UV laser cutting. Kapton rings were adhered to each end loop to reinforce the attachment point and focus deformation on the central strip. C) Image shows an assembled resistance band resting on a fingertip. Spring behavior was characterized using a custom tension testing machine. The plot shows two lines fitted to raw data from measurements of four resistance bands, used to approximate low-strain (yellow line) and high-strain (black line) stiffnesses. Scale bar: 5 mm.

Beginning with a sheet of elastomer film allows for simple control over spring length and width, important for determining ultimate spring stiffness. We cut resistance band profiles using a high-resolution UV laser engraver. We added loop features at each end to simplify system assembly, allowing the resistance band to be anchored around a post without the need for adhesives or bonding methods (Fig. 2B). To eliminate stress concentrations at the end loops, we lasercut 25 μm-thick polyimide washer-like discs and mechanically anchored them to the end loops with silicone epoxy. We validated spring stiffnesses with a custom tensile testing apparatus, described previously (Fig. 2C)^14^. We found that our particular design had a low strain stiffness of approximately 0.33 N/m up to a strain of 160%, within our 0.5 N/m stiffness design target. Even with manufacturing defects introduced during lasercutting and handling, our resistance bands could withstand strains greater than 300%. We believe this simple resistance band pre-tensioning design could be useful for other biohybrid mechanisms requiring light preload or restorative forces, saving physical space constraints and manufacturing complexity.

### Design and fabrication of millimeter-scale bistable flexures

Small, bistable flexures are conventionally monolithic: they comprise curved, rigidly clamped beams that are pushed in the center towards a critical point, until the beam quickly snaps through and settles in equilibrium on the opposing side^21^. Bistable snap-through behavior occurs in flexures via beam buckling, dynamically storing bending and compressive energies as the center displaces, manifesting as the biphasic force-displacement profile that captures bistable behavior.

Four-bar bistable flexures are usually made from rigid plastic, metal, or silicon-based materials that have elastic moduli above 1 GPa, requiring extreme beam aspect ratios to avoid yield strains at their ends. As a result, the majority of mm- and μm-scale bistable flexure literature reports stiffnesses greater than 1 N/m^22–25^. We found that employing the use of elastomers and mesoscale manufacturing methods again provided a pathway to achieve low actuation stiffnesses. We first used two-photon lithography (2PP) to generate replicas of desired flexure geometries with a rigid methacrylate-based resin. 2PP allowed us to precisely create a millimeter-scale component having sub-millimeter features including beam dimensions of 0.1 mm width, 1 mm length, and 0.7 mm depth. We then cast polydimethylsiloxane (PDMS) on these printed flexure replicas to make negative molds. Silane treatment of the PDMS molds minimized adhesion to the final cast soft PDMS flexure.

Flexures composed of four flat beams connected to a central element and anchoring points are not inherently capable of bistable behavior. Instead, we engineered bistability in such flexures by buckling the structure, pushing the anchor points inwards to a fixed offset position that raised the center to a new equilibrium position, thus allowing reversible push and snap through towards the opposite side (Fig. 3B)^26^. Experimental observations of our flexure show that near the critical point, beams exhibit a mirror symmetry across the horizontal axis: both pairs of beams bend outwards, as shown in Fig. 3B, step 3.

**Figure 3:**
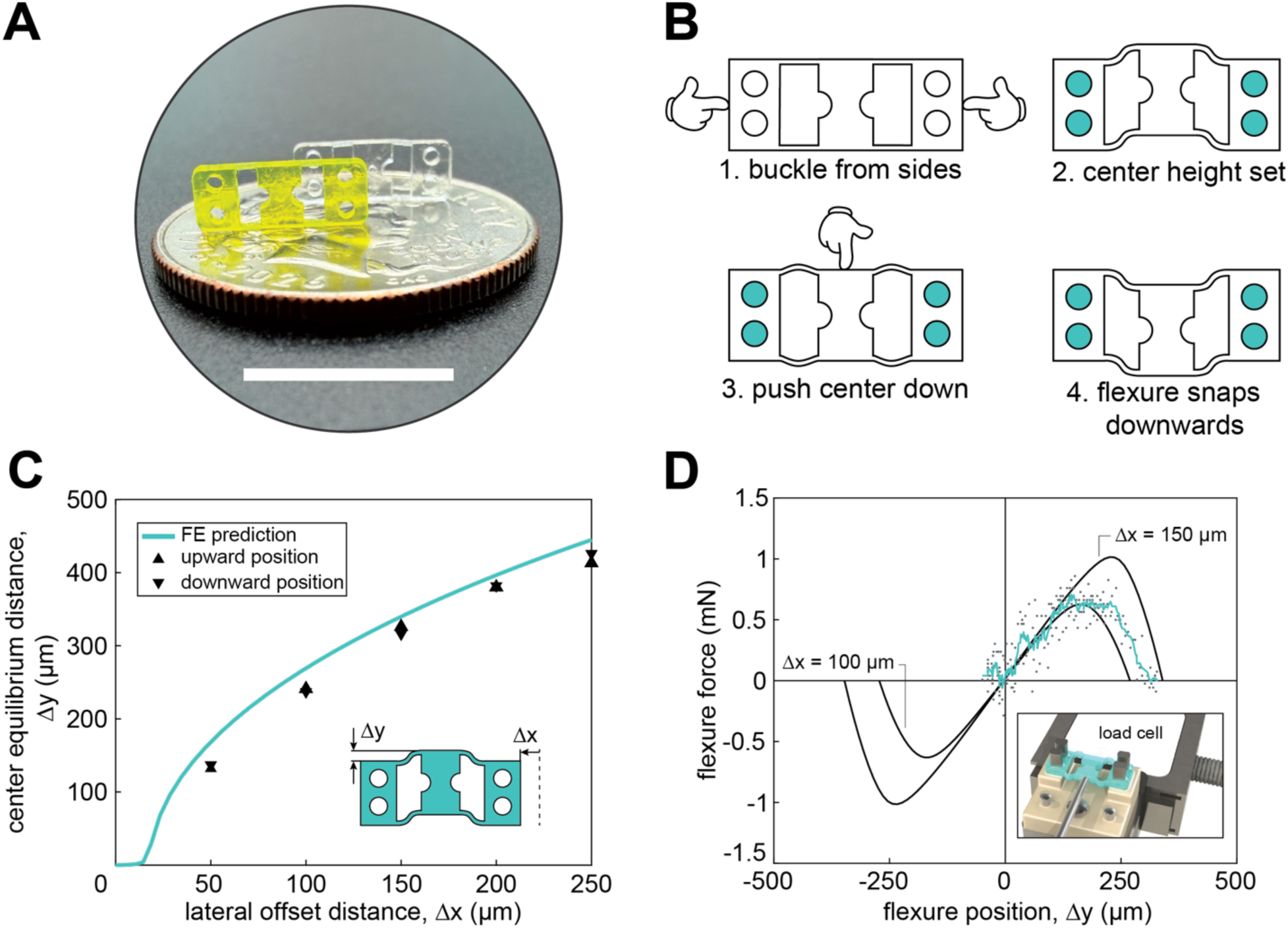
Mesoscale, soft bistable flexure. A) We used 2-photon lithography to 3D print the flexure mold positive (yellow), followed by casting to obtain the PDMS flexure structure (clear). Scale bar: 10 mm. B) We induce bistability by laterally buckling the four flexure beams such that each beam forces the center upwards (1). After fixing the ends in a shortened state (2), pushing the center past the midline (3) causes the flexure to flip towards the other stable position (4). C) We validated our design by matching finite element predictions of offset equilibrium position with experimental stereoscope measurements. D) Finite element predictions of reaction force were compared to measurements probing pre-buckled flexures past snap through on a custom tension testing machine, shown by a rendering in the bottom righthand corner. The teal curve was generated from compiling four measurements of a single flexure and using a moving average function (movmean, window length = 5, MATLAB R2024a). The experimental tail shown for Δy < 0 μm indicates that the flexure flipped downwards and lost contact with the probe. The solid lines are taken from FE simulations at offset distances of 100 μm and 150 μm, to show good agreement with experimental deviations from our offset distance of 125 μm.

This “pre-buckling” bistable design strategy has two advantages. First, force-displacement profiles can be tuned *in situ* by adjusting the distance of each end, helpful to account for biological variance in muscle output. Second, pre-buckled flexures have symmetric force-displacement profiles that can balance the load on antagonist muscle contraction, because the initial stresses generated from the offset reflect equally about a central critical point. This is unlike flexures that are fabricated to begin with a raised center, where pushing the center down produces a bending stress that grows monotonically, often delaying the critical point until passing the vertical midpoint of the flexure^24,25,27^. Testing upward and downward displacements of our flexure, we find nearly equidistant flipped positions and good agreement with finite element simulations of offset-center displacement curves (Fig. 3C, Supplemental Movie 1)^26^. Lastly, we measured flexure forces to confirm that stiffnesses were compatible with muscle contractile forces. Our experiments with a test flexure at an offset of 125 μm showed a peak force of approximately 0.6 mN, close to our design target (Fig. 3D). Moreover, our flexure configuration demonstrates a smooth transition from positive to negative stiffness, compared to the sharper force peaks of conventional bistable flexure designs.

### Assembly and bistable actuation of a biohybrid toggle switch

The simplest biohybrid bistable mechanism is a toggle switch that can be set and reset by the antagonistic contractions of muscle actuators, managing passive muscle tension and controlling flexure offset position. To fabricate the muscle actuators, we used our previously established protocols for producing contractile skeletal muscle from optogenetic C2C12 mouse myoblasts expressing a 470 nm blue light-sensitive ion channel that enables non-invasive control of contractility^16^. Myoblasts were mixed in a fibrin/Matrigel hydrogel solution, cast in ring-shaped injection molds, maintained in growth media containing fetal bovine serum for 2 days, and differentiated in media containing horse serum and human insulin-like growth factor 1 for 10 days to mature into functional actuators. The ring shape of our muscles enabled modular reuse of fabricated devices over multiple experiments. During differentiation, tissue constructs were exercised via optical stimulation for 15 minutes daily to further encourage contractility and hypertrophy.

Our design choices fit these system functionalities into a device of similar footprint to the muscle actuators to maximize the toggle switch packing density. We integrate the muscle, resistance bands, and bistable flexure with a 3D-printed, mm-scale linear motion system attached to a polycarbonate frame (Fig. 4A). The open space cut into the middle frame allowed us to place functional components above and below the frame, ensuring that each of the three functional components could apply forces close to a single plane, avoiding off-axis deflections. We included additional functionalities to enable fine tuning of muscle position, spring tension, and bistable stroke length, thus accounting for biotic and abiotic manufacturing variance. Specifically, to perform on-device tuning of bistable flexure equilibrium position, we mounted each end of the flexure on two 3D-printed, millimeter-scale double parallelogram flexures (Fig. 4A). Slotted cams located next to the outside edge of each flexure precisely moved the flexures inward to create the bistable offset, converting manual rotations into robust sub-mm position adjustment.

**Figure 4:**
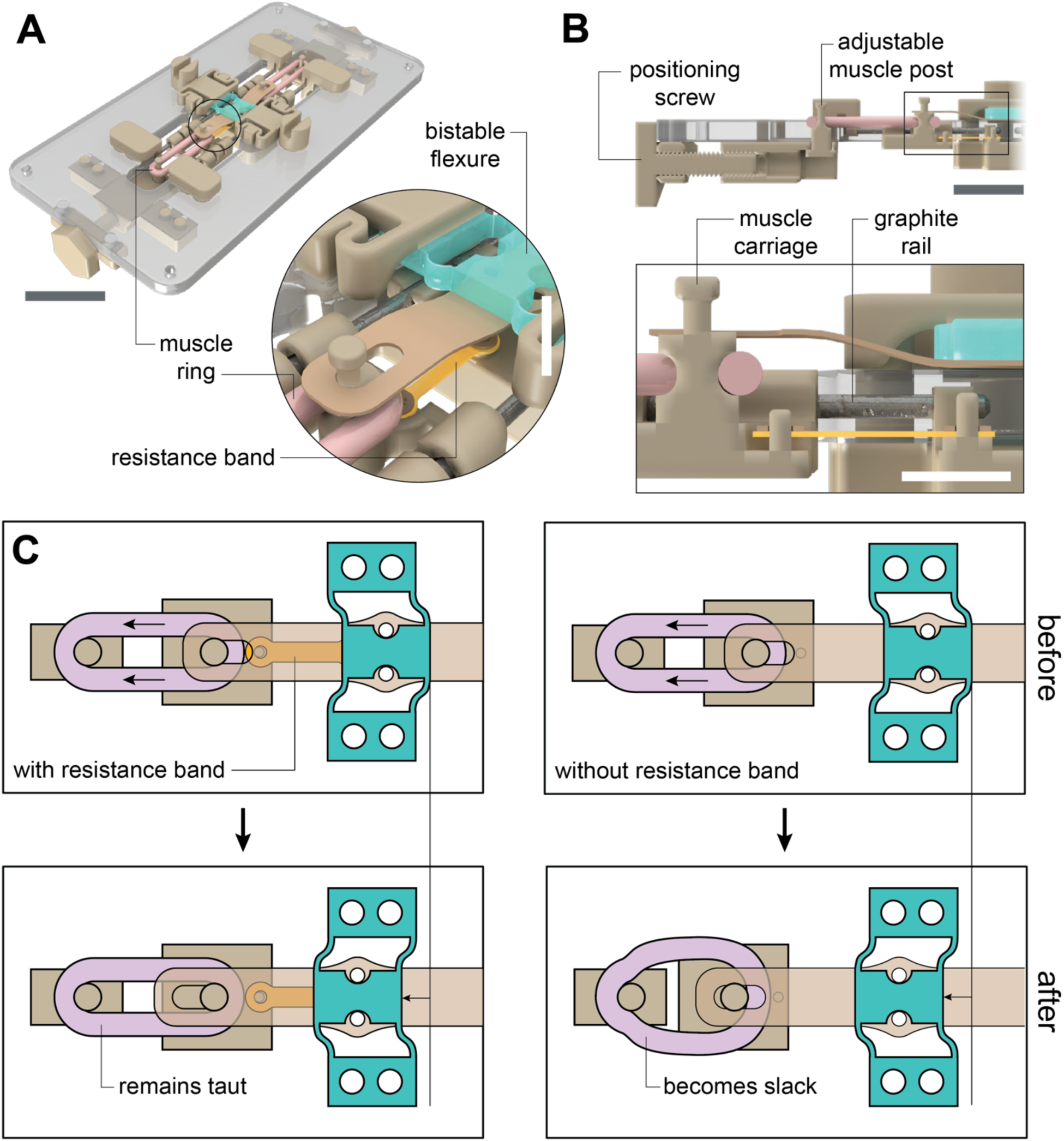
Biohybrid toggle switches. A) We mated 3D printed components with a polycarbonate frame to integrate all functional components into a single device. The system structure includes mating posts for the resistance bands, a shuttle to connect the muscle, band, and flexure, and a sliding back post to tune muscle tension. Compliant bistable flexure mounts allow for offset tuning via an integrated cam. Scale bars (left to right): 10 mm, 2.5 mm. B) A side view of the system shows how the components are arranged in-line. The carriage is mounted on graphite (pencil lead) linear rails due to their low friction, precision, and rigidity. Scale bars (top to bottom): 5 mm, 2.5 mm. C) Including a resistance band in the muscle carriage maintains muscle tension when the bistable flexure flips towards it. Otherwise, the muscle becomes slack, losing homeostatic loading and risking detachment from the posts. The resistance band also guarantees that the shuttle disengages from the bistable flexure, allowing the antagonistic muscle to actuate the flexure without interference.

The flexure-facing end, or front end, of each muscle was wrapped around a post connected to a millimeter-scale carriage which slid on 0.5 mm-diameter rails to constrain motions to one axis (Fig. 4B). The carriage transmitted muscle forces to the bistable flexure through a polyimide tab and also transmitted tensioning forces from the resistance band. The tab included a slot that allowed the carriage and flexure to be repeatably disengaged (Fig. 4C). The back end of each muscle ring connected to a post that can be positioned through rotations of a miniature lead screw having 00-80 thread geometry (corresponding to a pitch of 318 μm/rev), thus allowing for fine manual tuning during experiments. Following device assembly, we tensioned the muscles to engage them with the bistable flexure tabs (Fig. 5A).

**Figure 5:**
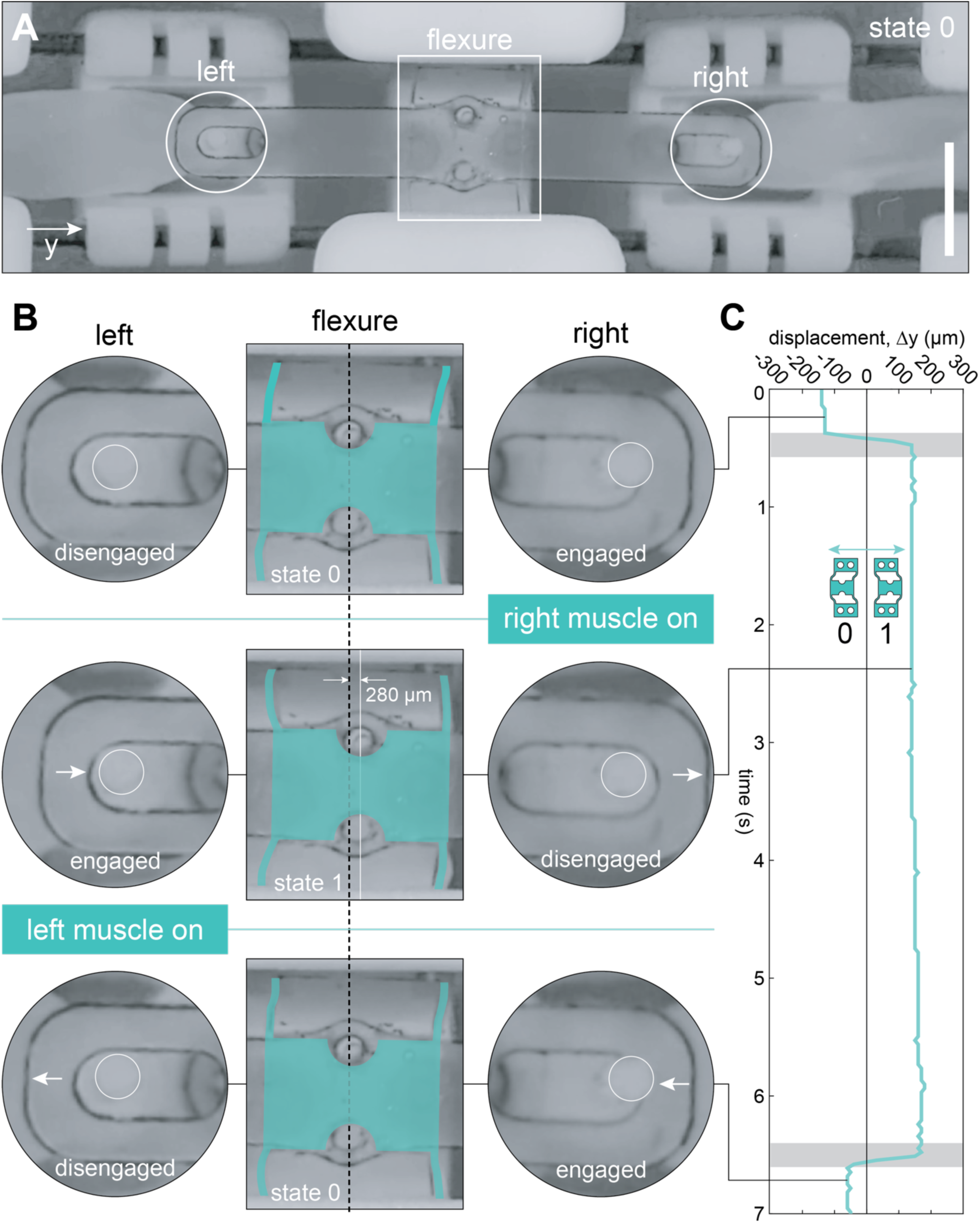
Muscle-powered bistable actuation. A) Mounting antagonistic muscle actuators on our mm-scale device, optical stimulation of the muscle enables successful back and forth snap-through of the flexure. Scale bar: 3 mm. B) Following from the initial “state 0” position, optical activation of the right muscle pulls the flexure and tab rightwards to state 1, causing flexure disengagement from the right carriage and engagement with the left carriage. The device reverses back to state 0 when the left muscle is activated. C) Plotting flexure position over time, we demonstrated that muscle-powered switching produces flexure states that persist without sustained actuation inputs. We oriented displacement values to be zero at the flexure’s center, the snap-through transition point. The grey shaded regions represent optical stimulation.

To demonstrate reversible muscle-driven actuation between bistable states, we stimulated the bottom muscle with a single light pulse (intensity of 50 μW/mm^2^), causing snap through from the upward “1” position to a downward “0” position (Fig. 5B). After snap-through, the actuating carriage post can be seen disengaging from the flexure tab as the opposing carriage engaged with the opposite tab. We then stimulated the top muscle in the same manner to switch from the “0” position back to the “1” position. During both movements, muscle activation actuated the bistable flexure 140 μm to its critical point, flipping an equal distance downward (Fig. 5C). The full snap-through motion occurs in 130 ms, giving an average center velocity of 2 mm/s. This implies an upper-limit switching frequency of 7.7 Hz.

## Discussion

State-of-the-art biohybrid robots have, to date, featured muscle constructs coupled to a spring-like skeleton to convert repeated contractions into a desired rhythmic motion such as locomotion or pumping. However, directly coupling muscles to skeletons intended to provide pre-tension and generate work constrains the ultimate work output and also limits the range of applications biohybrid actuators can serve. This work details a mechanism that expands the capability of engineered muscle actuators to perform mechanical switching operations. We integrate the muscle with a bistable flexure that structurally maintains equilibrium position, consuming no power after switching has occurred. With a single muscle attached, these flexures can be designed to amplify muscle displacement and power outputs. Here, we operate the flexure with an antagonistic muscle pair, showcasing a design concept that can apply towards several binary robotic applications such as payload release for implantable drug delivery devices and low-power soft gripping.

Core to the function of our biohybrid toggle switch is the design choice to decouple passive muscle tensioning from active operation of the bistable flexure. By tensioning muscle with a separate elastomeric resistance band that maintains a taut muscle position regardless of the position of the bistable flexure, we maintain optimal muscle tension in both stable states. Moreover, both the resistance band and the bistable flexure were designed to operate within the force-displacement envelope of our engineered C2C12 muscle actuators. Decoupling design requirements in this manner allowed us to independently tune bistable force-displacement profiles, muscle position, and muscle tension.

As with transistors, valves, and other switching components, miniaturization is essential to increase the utility of this actuation technology. A significant portion of our current device’s footprint is devoted to the carriage and linear rails. By leaving a small gap between the carriage sliding bearings and linear rail, we traded carriage alignment for lower parasitic frictions, which lowered the force transmission to the bistable flexure^19^. Designing low-stiffness flexures that minimize parasitic losses can avoid friction and maintain better muscle/mechanism alignment.

Despite these opportunities for further improvement, our study has established the first proof of principle that muscle actuators can be used to actuate bistable mechanisms, broadening the future application space for biohybrid machines.

## Materials and methods

### Passive tension muscle skeleton fabrication

We fabricated elastomer skeletons by casting elastomers in 3D-printed molds. To create the skeleton mold, we first designed and 3D printed a mold positive (Precision Model Resin, Form 4B, Formlabs). Positives were then washed with isopropyl alcohol (IPA) and let pieces rest overnight in an oven at 70 °C. Mold adhesion was reduced by conformally coating surfaces with vacuum-assisted silane vapor deposition (Trichloro(1H,1H,2H,2H-perfluorooctyl)silane, Sigma Aldrich) for four hours. Molds were casted with polydimethylsiloxane (PDMS, Sylgard 184, Dow) at a crosslinker-monomer ratio of 1:10 around each piece curing overnight at 70°C. After removing 3D-printed positives, the molds were silanized and then filled with PDMS to cast the skeletons. Lastly, skeletons were adhered to culture plates on the day of muscle mounting using silicone epoxy (QWIK-SIL, World Precision Instrument).

### Muscle ring mold fabrication

Muscle ring molds were cast from custom designed and 3D-printed mold masters (PLA, Prusa i3 MK3S+). After prints were cleaned in IPA, molds were cast in PDMS, placed in a vacuum chamber and degassed to remove bubbles, and cured overnight in an oven at 70 °C. Then, molds were peeled from the master molds and sterilized through autoclaving. Sterilized molds were stored in sealed containers until use. Directly before use, molds were washed with 1x DPBS (Gibco).

### Engineered muscle biofabrication and culture

We fabricated skeletal muscle actuators following our previously described 3D tissue culture protocols^14,16,19,28^, with the modification of adding E-64 to media. Optogenetically modified C2C12 myoblasts ChR2[H134R] were first cultured in growth medium (GM) composed of DMEM with additions of 4.5g/L of glucose, L-glutamine, and sodium pyruvate (Corning) supplemented with 10% (v/v) FBS (Life Technologies), 1% (v/v) penicillin/streptomycin (Fisher Scientific), and 1% (v/v) L-glutamine (Fisher Scientific).

One day before muscle ring fabrication, high-concentration Matrigel (Matrigel Basement Membrane Matrix, Corning) was thawed at 4 °C. On the day of muscle ring fabrication, thrombin (Sigma Aldrich) stock solution (100U/mL in 0.1% wt/v bovine serum albumin (BSA), Sigma Aldrich) was first set to thaw and supplemented growth media containing aminocaproic acid (6-Aminocaproic acid, Sigma Aldrich) stock solution (50mg/mL in DI) for a final concentration of 1mg/mL and 250µL of E-64 solution (E-64, Sigma Aldrich, 1mg/2.8mL) were put in a sterile ice bath. Prepared ring molds were then submerged in 1% w/v Pluronic (Pluronic F-127, Sigma Aldrich) in 1x DPBS for at least one hour.

After reaching at least 70% confluency, myoblasts were trypsinized (TrypLE Express, Gibco), harvested and counted into 3 million cell aliquots. Fibrinogen (Sigma Aldrich) was dissolved in growth media at a concentration of 8mg/mL and vortexed thoroughly, then set on ice. Cell aliquots were centrifuged at 1000 rpm for 5 minutes at room temperature into cell pellets. Cells were then resuspended in 59 μL of growth media, we added 1 µL of thrombin stock solution, 60 µL of Matrigel, and 180 µL of fibrinogen stock solution to the cell suspension in order. Molds were filled each with 120 μL of cell suspension, cured in an incubator for one hour, and submerged in GM.

After cells sufficiently compacted into a ring, we manually mounted them onto PDMS skeletons by wrapping each end around a pillar, allowing them to continue compaction for one day in GM. We then changed the tissue to differentiation medium (DM) containing DMEM with 4.5g/L of glucose, L-glutamine, and sodium pyruvate, supplemented with 10% (v/v) Horse Serum (Gibco), 1% (v/v) penicillin/streptomycin, 1% (v/v) L-glutamine, 1mg/mL of aminocaproic acid, 5ng/mL of IGF-1 (Sigma Aldritch), and 250 µL of E-64 stock solution. Maturing muscle actuators were maintained on DM for the remainder of tissue culture.

To promote muscle fiber alignment and development, we optically stimulated tissues daily with collimated LED light (470 nm, Thorlabs) for 15 minutes using a 2 Hz square wave and 20% duty cycle. This training was performed inside a biosafety cabinet, and differentiation media was changed after the stimulation period. Actuators were trained for at least 10 days before use in device experiments.

### Millimeter-scale resistance band fabrication

We fabricated passive-tensioning resistance bands through a combination of spin-coating and lasercutting manufacturing. First, elastomer sheets were prepared by spin coating silicone prepolymer onto acrylic substrates. Acrylic disc patterns (3mm-thick) were lasercut (Fusion M2, Epilog Laser) from stock material, cleaned with IPA, and mounted onto the spin coater vacuum chuck. Silicone prepolymer was prepared by mixing two-part elastomers (Ecoflex 00-10, Smooth-On, Inc.) and 10% w/w silicone thinner (Silicone Thinner, Smooth-On, Inc.) in a planetary centrifugal mixer (330-100 SE, FlackTek) at 2500 rpm for one minute. A dollop of silicone prepolymer was then placed on the acrylic substrate and spun at 3000 rpm for 45 seconds. Silicone-coated substrates were then removed and placed in an oven to cure at 70 °C overnight.

Resistance band spring patterns were patterned in arrays using a UV laser engraver (F2 Ultra, xTool), with respective speed, power, and frequency settings of 10 mm/s, 80%, and 150 kHz. Similarly, 0.25 mm-thick polyimide sheets taped to an acrylic substrate were laser cut into washer shapes. The polyimide reinforcement rings were pasted to the resistance bands with silicone epoxy (Sil-Poxy, Smooth-On, Inc.), ensuring sufficient epoxy covered the surface. After curing, a 0.35 mm-diameter punch cleared excess epoxy from the center of the disc and formed mounting holes. Springs were then peeled from the substrate and stored on a miniature rack until use.

### Resistance band validation

Resistance bands were characterized for film thickness and force output to check fabricated results against design targets. Films were characterized by cutting a portion of film out and measuring the difference in height from the substrate to the top of the film. A laser microscope was used (VK-X1000, KEYENCE) to measure 5 regions along each spin coated sample (VK-H1XME Multi-file Analysis Application, KEYENCE), reporting average values in Fig. 2C. Force measurements were conducted with a custom tensile testing apparatus^14^ using an off-the-shelf force sensor (LSB200, 10 g, FUTEK) and amplifier (IAA100, FUTEK).

### Bistable flexure fabrication

To fabricate the soft flexures, mold positives were first 3D-printed using two-photon photolithography (5x objective, UpPhoto, UpNano). Positives were placed on a flat PDMS substrate cured in a petri dish and pressed to remove any air gaps between surfaces. The positives and substrate were then silanized and cured at 70 °C overnight. Pouring 1:10 PDMS prepolymer into the petri dish formed the mold negative. Care was taken to remove any small bubbles trapped in the corners and holes of the mold positives, using a small wire. After curing at 70 °C, the mold was removed from the petri dish and silanized. Then, 1:10 PDMS prepolymer was slowly added to the mold to form the final flexure components, and components were removed by tweezer after curing.

### Bistable flexure validation

We conducted finite element (FE) modeling, manufacturing validation, and force measurement experiments to validate the bistable flexure fabrication process. Flexure offset positioning and force-displacement relationships were modeled in ABAQUS using a static solver and a CPS4R mesh. We chose 1 MPa as an approximate linear elastic modulus for PDMS. To validate the offset-center displacement prediction shown in Fig. 3C, sample flexures were pinned to custom steel fixture mounted on a manual precision linear stage (MS1S/M, Thorlabs) to control the offset distance. The flexure was manually flipped back and forth, and images showing center position were taken using an overhead stereo microscope (Stemi 305, Zeiss). To experimentally validate force-displacement relationships, flexure samples were pinned to a mount connected to a custom tensile-testing apparatus and a steel probe slowly pushed the center downwards until snap-through occurred.

### Device fabrication and assembly

The actuation system was fabricated by press fitting 3D-printed mm-scale components into a polycarbonate frame. The frame was machined on a 3-axis milling machine (ProtoTrak) using 2D pocket and drilling toolpaths (Fusion, Autodesk). System components were 3D printed using stereolithography (Precision Model Resin, Form 4B, Formlabs) and washed. To suppress excess material at the component bases, the early-layer exposure and offset settings were modified in the printing software (PreForm, Formlabs). At a print height of 0 mm, the exposure dose was set to 140 mJ/cm² and the offset to 0.300 mm; at a print height of 0.600 mm, the exposure dose was set to 19 mJ/cm² and the offset to 0.008 mm. Irradiance was set to 13 mW/cm² for both settings. The printed components were bonded to one another and to the frame using a UV-curing adhesive (Norland, NOA 61). Components were bonded to each other and the frame with light-curing adhesive. The flexure was bonded to the polyimide film using silicone epoxy (Sil-Poxy, Smooth-On). Graphite rails made by cutting mechanical pencil leads (0.5 mm Hi-Polymer Super Lead, Pentel) were snap fit into rail blocks and held by friction. Flexures were then adhered to mounts with silicone epoxy. Lastly, resistance bands were looped around posts on the muscle carriage and central module by tweezer. After assembly, devices were washed in IPA, UV sterilized, and stored in airtight containers until use.

### Bistable actuation demonstration and analysis

During experimental demonstrations, devices were submerged in DMEM in a deep-well petri dish and muscle actuators were wrapped around posts with the aid of a stereoscope. Muscles were then tensioned and aligned with the pre-buckled flexure so that each stage was minimally engaged when loaded by the flexure. Controlled by a 1 Hz square wave and 20% duty cycle, collimated light optically stimulated cells to trigger contractions and toggle switching. An iPhone 17 connected to a stereo microscope filmed toggle switch operation. Shuttle post and flexure center positions were tracked using ImageJ (Manual Tracking, Fiji).

## Acknowledgements

The authors acknowledge Mark Belanger and the Edgerton student shop for guidance on metrology and frame machining, the MIT Central Machine Shop for fixture fabrication services, the labs of Profs. Xuanhe Zhao and John Hart for equipment access, and Prof. Martin Culpepper for helpful discussions. The authors acknowledge funding support from the Army Research Office Early Career Program and PECASE Award (W911NF2210126, awarded to Ritu Raman) and the MIT Engineering Excellence Postdoctoral Fellowship (awarded to Ronald Heisser).

